# Structural basis for activation of Dot1L methyltransferase on the nucleosome by histone H2BK120 ubiquitylation

**DOI:** 10.1101/503128

**Authors:** Cathy J. Anderson, Matthew R. Baird, Allen Hsu, Emily H. Barbour, Yuka Koyama, Mario J. Borgnia, Robert K. McGinty

**Affiliations:** Department of Biochemistry and Biophysics, School of Medicine, The University of North Carolina, Chapel Hill, NC 27599, USA; Division of Chemical Biology and Medicinal Chemistry, Eshelman School of Pharmacy, The University of North Carolina, Chapel Hill, NC 27599, USA; Genome Integrity and Structural Biology Laboratory, National Institute of Environmental Health Sciences, National Institutes of Health, Department of Health and Human Services, Research Triangle Park, NC 27709; Present address: Department of Cell Biology, Harvard Medical School, Boston, MA 02115, USA

**Keywords:** chromatin, structural biology, single particle, cryo-EM, histone methyltransferase, nucleosome, Dot1L

## Abstract

Histone H3 lysine 79 (H3K79) methylation is enriched on actively transcribed genes, and its misregulation is a hallmark of leukemia. Methylation of H3K79, which resides on the structured disk face of the nucleosome, is mediated by the Dot1L methyltransferase. Dot1L activity is part of a trans-histone crosstalk pathway, requiring prior histone H2B ubiquitylation of lysine 120 (H2BK120ub) for optimal activity. However, the molecular details describing both how Dot1L binds to the nucleosome and why Dot1L is activated by H2BK120 ubiquitylation are unknown. Here we present the cryo-EM structure of Dot1L bound to a nucleosome reconstituted with a site-specifically ubiquitylated H2BK120. The structure reveals that Dot1L engages the nucleosome acidic patch using an arginine anchor and occupies a conformation poised for methylation. Ubiquitin directly interacts with Dot1L and is positioned as a clamp on the nucleosome interacting region of Dot1L. Using our structure, we identify point mutations that disrupt the nucleosome-specific and ubiquitin-dependent activities of Dot1L. This study establishes a path to better understand Dot1L function in normal and leukemia cells.

## Introduction

Histone lysine methylation contributes to the regulation of transcription by tuning the recruitment of effector proteins to specific genomic sites (Hyun et al., 2017). It exists in mono-, di-, and trimethylated (me1, me2, and me3) forms, and functional outcomes depend on both the methylated histone residue and degree of methylation (Greer and Shi, 2012). Most well-characterized sites of histone lysine methylation are found in the flexible N-terminal tails of histones (Zhao and Garcia, 2015). One counterexample is histone H3 Lys79 (H3K79), which is solvent exposed on the structured disk face of the nucleosome (Luger et al., 1997). H3K79 methylation is observed within transcriptionally active genes, and methylation levels are highly correlated with gene expression (Schübeler et al., 2004; Wang et al., 2008; Wood et al., 2018). In human cells, H3K79me2/me3 are enriched immediately after transcription start sites and decrease gradually across gene bodies, while H3K79me1 is distributed more broadly across the bodies of active genes (Wang et al., 2008).

Dot1L/KMT4 (disruptor of telomeric silencing-1 like/lysine methyltransferase 4) is the primary H3K79 methyltransferase in human cells and is conserved across eukaryotes (Feng et al., 2002; Lacoste et al., 2002; Ng et al., 2002a; van Leeuwen et al., 2002). Rather than having the characteristic SET (Su(var)3–9, Enhancer-of-zeste, Trithorax) domain found in other histone lysine methyltransferases (Dillon et al., 2005), Dot1 proteins have a catalytic domain resembling class I methyltransferase domains found in DNA and protein arginine methyltransferases (Min et al., 2003; Sawada et al., 2004). While known to participate in several transcriptional elongation complexes (Wood et al., 2018), Dot1L can bind to and methylate H3K79 in nucleosomes in isolation (Feng et al., 2002; Min et al., 2003). Histone H3 alone is a poor substrate for Dot1L, suggesting that Dot1L requires non-H3 surfaces of the nucleosome for substrate binding and/or activity (Feng et al., 2002; Lacoste et al., 2002; Ng et al., 2002a).

Efficient methylation of H3K79 in cells requires prior ubiquitylation of H2BK120 (Briggs et al., 2002; Kim et al., 2005; Ng et al., 2002b). H3K79me2/me3 are significantly decreased without change to H3K79me1 following knockdown of the H2BK120-targeting ubiquitin E3 ligase, Bre1, in human cells or upon mutation of H2BK120 in *S. cerevisiae* (Kim et al., 2005; Shahbazian et al., 2005). Using designer nucleosomes assembled with monoubiquitylated H2BK120 (H2BK120ub), this trans-histone crosstalk between H2BK120ub and H3K79 methylation has been shown to be direct and require only the catalytic domain of Dot1L (McGinty et al., 2008). Previous studies implicate the C-terminal tail of ubiquitin and the N-terminal tail of histone H2A in mediating ubiquitin-dependent Dot1L activity (Holt et al., 2015; Zhou et al., 2016). The N-terminal tail of H4 has also been shown to be important for Dot1L activity independent of H2B ubiquitylation (Fingerman et al., 2007; McGinty et al., 2009). In recent years, Dot1L has emerged as a potential therapeutic target for MLL-rearranged leukemias because the catalytic activity of Dot1L is required for leukemogenic transformation following MLL-fusion translocations (Bernt et al., 2011; Winters and Bernt, 2017). Yet, key molecular details describing how Dot1L binds to the nucleosome and is activated by H2B ubiquitylation remain elusive.

Here we report the 3.9 Å cryo-EM structure of the methyltransferase domain of human Dot1L bound to a designer nucleosome core particle assembled with a site-specifically modified H2BK120ub (Figure 1A). The structure captures Dot1L in a poised state in which Dot1L uses an arginine to anchor to the acidic patch on the nucleosome and is clamped in place by ubiquitin. In the structure, the catalytic site of Dot1L is separated from H3K79, indicating that Dot1L and/or the nucleosome must undergo conformational rearrangement from a poised to an active state for methylation.

**Figure 1.**
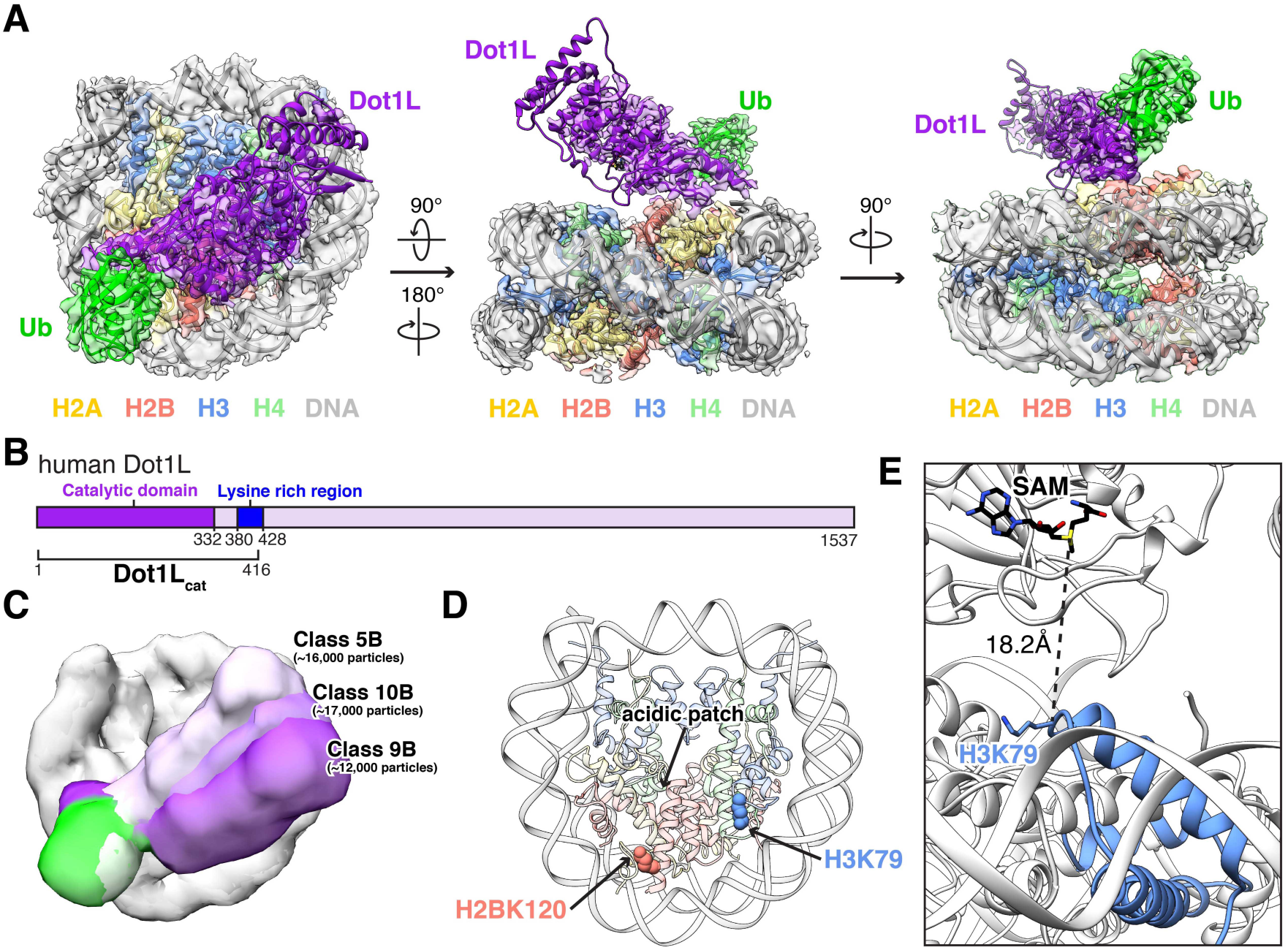
Cryo-EM structure of Dot1L-H2BK120ub nucleosome complex. **A**, Three orthogonal views of the complex with reconstructed map overlaying molecular model, colored as indicated. **B**, Scheme of human Dot1L highlighting the catalytic domain (dark purple), the lysine rich region (blue), and the minimal Dot1L_cat_ fragment used in our studies. **C**, Overlay of 3D subclasses showing conformational heterogeneity of Dot1L (shades of purple) and ubiquitin (shades of green). **D**, Nucleosome structure with acidic patch and sites of ubiquitylation (H2BK120) and Dot1L-mediated methylation (H3K79) indicated. **E**, Zoomed view of Dot1L active site and targeted lysine in our model with distance between H3K79 Cα and SAM cofactor methyl donor measured.

## Results

### Cryo-EM structure of Dot1L bound to H2BK120ub nucleosome

In order to visualize the molecular details of Dot1L function on nucleosomes and activation by H2BK120 ubiquitylation, we first needed to prepare homogeneously site-specifically ubiquitylated nucleosomes. To this end, we generated and purified an H2BK120ub analog by crosslinking ubiquitin with a Gly76Cys mutation at its terminal residue to an H2BK120C mutant using dichloroacetone (DCA) as previously described (Figure S1) (Morgan et al., 2016). Importantly, we observe slightly increased Dot1L methyltransferase activity when using nucleosomes reconstituted with two copies of this purified DCA-crosslinked analog as compared to nucleosomes assembled with two copies of a more precise semisynthetic analog of H2BK120ub containing a single Gly76Ala point mutation in ubiquitin (McGinty et al., 2009). As this Gly76Ala analog was previously established to be indistinguishable from natively linked H2BK120ub in a Dot1L methyltransferase assay, we can extrapolate that Dot1L is activated by our DCA-crosslinked analog similarly to, if not to a greater extent than native H2BK120ub. We reconstituted our homogeneous DCA-crosslinked H2BK120ub nucleosome core particles with a minimal catalytic fragment of Dot1L (Dot1L_cat_) (Figure S1). This fragment spanning amino acids 2–416 includes the catalytic domain of Dot1L and an adjacent lysine-rich sequence that was previously identified to be necessary and sufficient for nucleosome binding and activity (Figure 1B) (Min et al., 2003). Although stable in solution, Dot1L_cat_ dissociated from H2BK120ub nucleosomes during cryo-EM sample preparation under all conditions tested. To improve complex stability, we crosslinked the complex with glutaraldehyde prior to vitrification. This allowed for visualization of a Dot1L_cat_-H2BK120ub nucleosome complex by cryo-EM.

We initially resolved a 1:1 Dot1L_cat_-H2BK120ub nucleosome complex from 3D reconstruction of ~100,000 particles at 3.5 Å resolution (Figure S2, Classes 3 and 4). In this reconstruction, high resolution features in the nucleosome region of the density map are consistent with the estimated resolution. However, conformational heterogeneity in Dot1L and ubiquitin limit resolution in their respective regions of the reconstructed map. To improve Dot1L and ubiquitin visualization, we reclassified the particles used for the 3.5 Å reconstruction into four subclasses, leading to the identification of two major conformational states (Classes 2A and 4A). We used Class 2A to generate a 3.9 Å reconstruction, that while having a lower overall resolution than our initial map, has improved density for Dot1L and ubiquitin generally, and does not compromise high resolution information at the Dot1L-nucleosome interface. The observed conformational heterogeneity in Dot1L and ubiquitin is further illustrated by finer classification into 10 subclasses, where a continuum of Dot1L and ubiquitin conformations is observed (Classes 5B, 9B, and 10B) (Figure 1C). As these classes did not have sufficient particles to allow for high resolution reconstruction, subsequent modeling was completed using the 3.9 Å map. It is worth noting that we anticipated a 2:1 complex (i.e. two copies of Dot1L per ubiquitylated nucleosome) due to the expected ability of the methyltransferase to bind to each face of the psuedosymmetric nucleosome core particle (McGinty et al., 2009; Zhou et al., 2016). However, we initially observed only sparse density in the regions corresponding to Dot1L and ubiquitin on one nucleosome face. This is likely due to loss of Dot1L on the other nucleosome face during sample freezing and the resulting conformational heterogeneity of ubiquitin when unconstrained by Dot1L. This is consistent with a previous crystal structure of an unbound H2BK120ub nucleosome that lacks defined density for ubiquitin (Machida et al., 2016). Only in the finer classification described above did we identify minor class averages with definitive density for a 2:1 Dot1L:nucleosome complex (Class 4B) or density for ubiquitylated nucleosomes only (Class 7B), demonstrating that despite crosslinking with glutaraldehyde our particles include an ensemble of Dot1L:nucleosome stoichiometries.

With the optimal 3.9 Å resolution reconstructed map in hand, we generated a Dot1L_cat_-H2BK120ub nucleosome model by docking high resolution crystal structures of the nucleosome core particle, one Dot1L catalytic domain, and one ubiquitin into the map followed by iterative manual model building and real-space refinement (Figure 1A). Although present on the opposite nucleosome faces, ubiquitin was not modeled on the nucleosome face not bound by Dot1L due to poorly resolved density. In our model, Dot1L anchors to the nucleosome on the edge of the nucleosome acidic patch – an emerging hot spot for nucleosome binding (Figure 1D). We do not observe density for the lysine rich region C-terminal to the catalytic domain, suggesting that it binds to the nucleosome in a heterogeneous manner. In fact, no density is unaccounted for by the model in this region or any other region of the map. Ubiquitin, which is covalently attached to H2BK120, occupies a position lifted off the nucleosome surface and packs against the nucleosome-binding region of Dot1L. Interestingly, Dot1L does not engage the nucleosome surface surrounding the targeted H3K79 residue. Rather, a gap exists between H3K79 and the Dot1L active site. Because we do not observe obvious density for the S-adenosylmethionine (SAM) cofactor in the Dot1L active site, we aligned our model with a Dot1L-SAM co-crystal structure, allowing us to determine the position of SAM. This alignment shows an 18 Å distance between the donor methyl carbon of SAM and the H3K79 Cα atom (Figure 1E). We hypothesize that our model represents a poised state and that conformational rearrangement of the nucleosome and/or reorientation of Dot1L relative to the nucleosome is required for catalysis to occur. Interestingly, mass spectrometric analysis of our reconstituted Dot1L_cat_-H2BK120ub(DCA) complex shows that the majority of H3 is monomethylated (Figure S1). As we did not supplement our complex with SAM, this observation suggests that Dot1L_cat_ purified with SAM bound and has subsequently methylated the nucleosome. Therefore, the majority of our reconstructed particles contain a bound Dot1L_cat_ that is poised for H3K79 dimethylation.

It is worth noting that the poised conformation of Dot1L in our cryo-EM model is similar to a molecular replacement model of a non-glutaraldehyde crosslinked, Dot1L-H2BK120ub nucleosome complex that we solved using an 8 Å resolution X-ray diffraction dataset (Dot1L Cα RMSD = 10.4 Å following alignment by histones) (Figure S3 and Table S1). This crystal structure shows a 2:1 Dot1L-H2BK120ub nucleosome complex in which Dot1L is bound to the identical surface of the nucleosome in a nearly identical manner, but held even farther away from H3K79 than in our cryo-EM model. We initially considered this poised conformation in our crystallographic model to be an artifact of crystal packing due to the presence of a symmetry-related copy of Dot1L wedged between the Dot1L catalytic site and its targeted nucleosome. However, the convergence of our models generated by cryo-EM and X-ray crystallography suggests a functional role for the observed conformation. Moreover, as our crystallographic complex was not glutaraldehyde crosslinked, it is unlikely that crosslinking has artificially forced Dot1L into the poised state observed in our cryo-EM structure. As the majority of particles in our cryo-EM dataset are classified into 3D classes with visible density for Dot1L in the poised conformation, we expect that this is the preferred equilibrium position of Dot1L on the nucleosome.

Significant variation of resolution exists across the reconstructed cryo-EM map (Figure S4). The nucleosome dominates particle alignments and is thus well-resolved allowing high confidence modeling of most histone side chains. Even though we initiated nucleosome modeling with a high resolution nucleosome crystal structure, the final model closely resembles a published cryo-EM structure of an unbound nucleosome (RMSD = 1.2 Å across shared Cα atoms), which diverges from crystal structures most notably in the slight splaying of the H2B αC helices away from the nucleosome surface (Bilokapic et al., 2018). In contrast to the nucleosome volume, the regions of the reconstructed map corresponding to Dot1L and ubiquitin exhibit more limited high resolution features. The catalytic domain of Dot1L was easily docked using a well-resolved nucleosome binding loop and surrounding features including the αK helix that closely borders the ubiquitin density. Secondary structure in nearby Dot1L regions including the catalytic site also aligns well to the map, but local resolution is limited due to the aforementioned conformational heterogeneity. More distant regions and large loops of Dot1L are less well resolved in our map. Ubiquitin was most challenging to model due to its smaller size and spheroidal shape. To improve rigid body docking of the ubiquitin structure, we masked the volumes of Dot1L and ubiquitin and continued refinement (Figure S5). While this approach failed to generate side chain resolution for Dot1L or ubiquitin, clear density for secondary structure elements allowed high confidence docking of ubiquitin and further confirmed Dot1L placement in the nucleosome binding and active site regions.

### Dot1L-nucleosome interaction

The catalytic domain of Dot1L binds to the nucleosome using a long pre-structured loop between Dot1L β9 and β10 sheets (Figure 2A). We clearly resolve main chain and select side chain density for this Dot1L nucleosome-interaction loop. The most striking feature within the loop is the Dot1L Arg282 side chain that clearly inserts into a cavity at the edge of the nucleosome acidic patch, in position to make a salt bridge and a hydrogen bond with the side chains of H2AE56 and H2BQ47, respectively. Two other residues in the loop, Arg278 and Asn280, are positioned to make direct nucleosome interactions, but their side chains are not clearly resolved in the map. To validate the importance of the nucleosome-interaction loop for Dot1L methyltransferase activity, we tested alanine mutants of key loop residues using an enzyme-coupled methyltransferase assay. We first established that the assay could detect Dot1L_cat_ activity on unmodified nucleosomes and nucleosomes containing a near-native semisynthetic H2BK120ub analog with a Gly76Ala mutation in ubiquitin and that the time points and enzyme concentrations we used were in the linear range of the assay (Figure 2B and Figure S1). We observe a ~6-fold enhancement of Dot1L_cat_ activity on H2BK120ub nucleosomes as compared to unmodified nucleosomes. This is a smaller enhancement than previously described using a direct assay with ^3^H-SAM and may result from intrinsic differences between the assays, but the same overall trend is observed (McGinty et al., 2009). Because Dot1L activity could be detected on both ubiquitylated and unmodified nucleosomes, we were able to assess ubiquitin-dependent and independent effects of Dot1L mutations. As predicted by our model, Dot1 Arg282Ala leads to a 90% reduction in H3K79 methylation on ubiquitylated nucleosomes and nearly abolishes all activity on unmodified nucleosomes. Asn280Ala and Arg278Ala mutations also result in losses of activity, especially on unmodified nucleosomes, although to a lesser degree than the Arg282Ala mutant Dot1L. Notably, the best resolved side chain in Dot1L at the nucleosome interface in our cryo-EM map is also the most important side chain for Dot1L activity regardless of H2B ubiquitylation. The Dot1L nucleosome interaction loop is well conserved in higher eukaryotes but divergent in *S. cerevisiae* ortholog Dot1p, leaving open the possibility that Dot1p binds nucleosomes and is activated by ubiquitin through a distinct mechanism compared to human Dot1L (Figure 2C).

**Figure 2.**
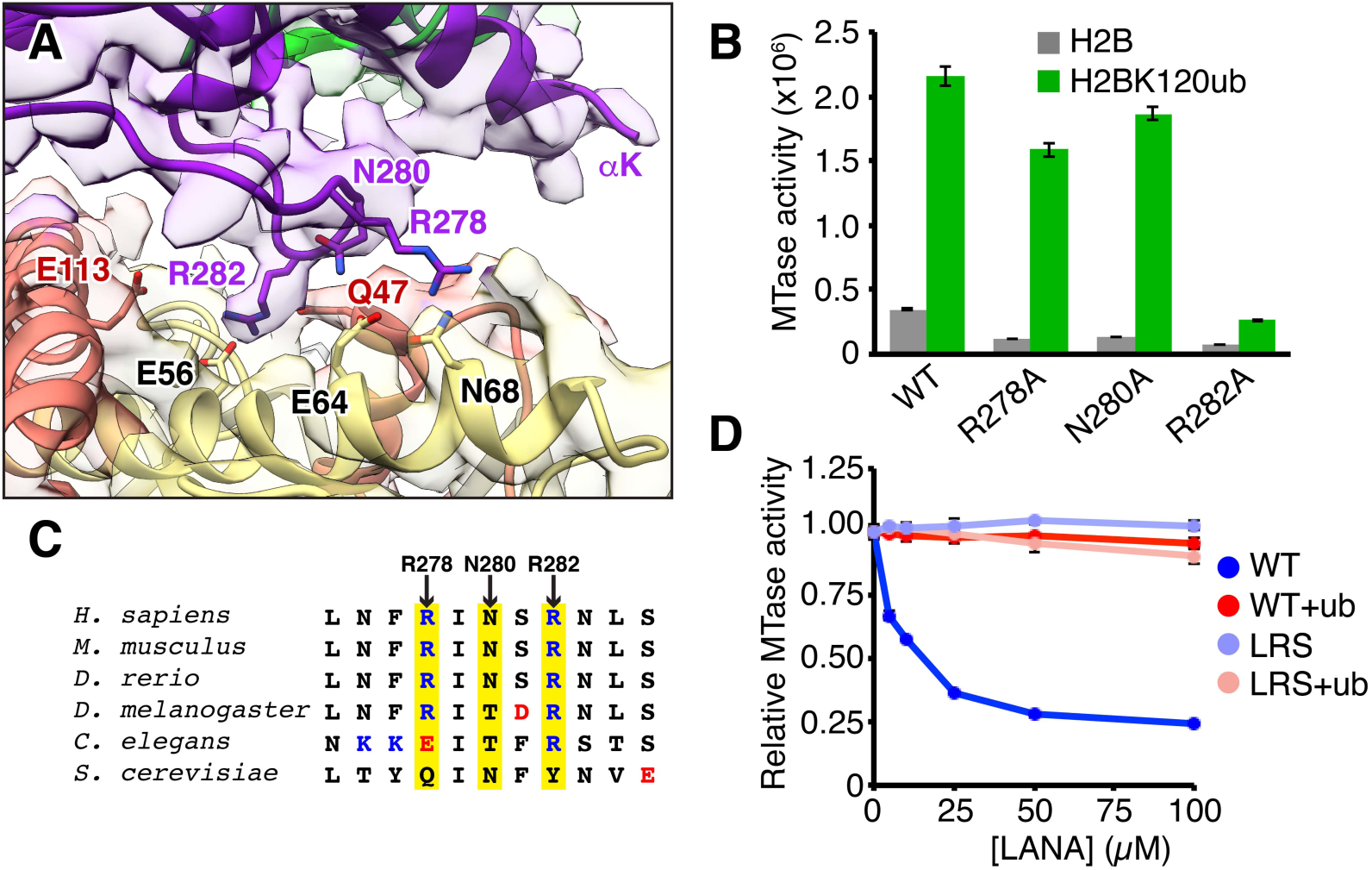
Dot1L interacts with the nucleosome acidic patch. **A**, Zoomed view of Dot1L nucleosome-interaction loop engaged with nucleosome acidic patch surface. Model, reconstructed map, and relevant side chains are shown. **B**, Quantified Dot1L methyltransferase assay using indicated mutants with unmodified (H2B, grey) and ubiquitylated (H2BK120ub, green) nucleosomes. **C**, Sequence alignment of Dot1L nucleosome-interaction loop with positively and negatively charged residues colored blue and red, respectively, and key positions highlighted in yellow. **D**, Competitive methyltransferase assay with varying concentrations of wild-type (WT) or nucleosome binding-deficient (LRS = LRS 8–10 AAA) LANA fusion proteins on unmodified (shades of blue) and H2BK120ub (+ub, shades of red) nucleosomes. Five replicates performed for all assays and means and standard deviations are shown.

While our structure shows that Dot1L binds to the acidic patch of ubiquitylated nucleosomes, an alternative binding mode has been suggested for Dot1L on unmodified nucleosomes (Zhou et al., 2016). To test whether Dot1L requires the acidic patch for activity on unmodified nucleosomes, we performed competitive methyltransferase assays by titrating a GST-fusion of the nucleosome acidic patch-binding 23 amino acid LANA (latency-associated nuclear antigen from Kaposi’s sarcoma-associated herpesvirus) sequence into our assays (Figure 2D). Wild-type LANA, but not a nucleosome binding-deficient mutant (residues 8–10, LRS, mutated to AAA), is able to inhibit Dot1L activity on unmodified nucleosomes in a dose dependent manner. While not ruling out alternative binding modes, this suggests that Dot1L activity on ubiquitylated and unmodified nucleosomes shares a similar acidic patch interaction dependence. To our surprise, LANA fails to substantially inhibit Dot1L enzymatic activity on ubiquitylated nucleosomes even at very high concentrations (Figure 2D). This is not due to the GST-fusion, as pre-assay cleavage of the GST tag leads to identical results (Figure S6). This observation could be secondary to tighter Dot1L binding to H2BK120ub nucleosomes or the inability of LANA to bind to H2BK120ub nucleosomes altogether. Further investigation shows that GST-LANA forms a stable complex with unmodified nucleosomes, but is unable to bind to H2BK120ub nucleosomes (Figure S6). Importantly, this presents a mechanism in which H2BK120 ubiquitylation may activate Dot1L in isolation, but may also further promote Dot1L activity in cells by acting as a gatekeeper to the acidic patch and neutralizing competition for nucleosome acidic patch binding.

### Dot1L-ubiquitin interaction

Ubiquitin occupies a position lifted off the nucleosome surface and rests against the αK helix of Dot1L that scaffolds the structure of the Dot1L nucleosome-interaction loop. Dot1L and ubiquitin exhibit conformational flexibility in our cryo-EM dataset. However, docking of Dot1L and ubiquitin into finely parsed 3D classes followed by alignment by Dot1L shows that Dot1L and ubiquitin move as a unit, suggesting a direct interaction (Figure S7). Previously reported comprehensive mutagenesis of the ubiquitin surface identified only two ubiquitin side chains necessary for Dot1L activation, Leu71 and Leu73, both located near the C-terminus of ubiquitin. Therefore, even though the ubiquitin-Dot1L interface appears large, we expected to find that only the small region of Dot1L adjacent to ubiquitin Leu71 and Leu73 would be responsible for ubiquitin-dependent activity. While the ubiquitin Leu71 side chain is poorly resolved in our map, Leu71 is at the end of the β4 strand that is part of the globular-fold of ubiquitin, so we can confidently place its side chain in proximity to Dot1L αK residues Leu322 and Phe326 (Figure 3A). We have less confidence in the location of ubiquitin Leu73 in our model as it resides in the flexible ubiquitin tail. However, based on its importance for ubiquitin-dependent Dot1L activity, we hypothesized that Leu73 makes direct contact with a hydrophobic surface on Dot1L. A likely surface is the exposed Ile290 side chain of Dot1L. We tested the effects of alanine mutations of Ile290, Leu322, and Phe326, as well as Glu323 and Lys330 that are also positioned at the Dot1L-ubiquitin interface but more distant from ubiquitin Leu71 and Leu73 (Figure 3B). To assess H2BK120ub-specific effects, we compared fold-increases in Dot1L activity due to H2B ubiquitylation for each of the Dot1L mutants. The Phe326Ala mutation effectively abolished all ubiquitin-dependent activity of Dot1L, with a calculated H2BK120ub fold-increase of 0.9 (compared to 6.2 for wild-type Dot1L). An Ile290Ala mutation also preferentially disrupted H2BK120-depedendent activity of Dot1L, with a calculated fold-increase of 2.8. A Dot1L Leu322Ala mutant methylated both unmodified and ubiquitylated nucleosomes with near wild-type activity, so we prepared a more severe Leu322Glu mutant Dot1L. This mutation resulted in a more pronounced reduction in activity on H2BK120ub nucleosomes than unmodified nucleosomes, leading to a ubiquitin fold-increase of 1.3. Glu323Ala and Lys330Ala mutations, which are not in the vicinity of ubiquitin Leu71 or Leu73 in our model, had minimal effects on Dot1L enzymatic activity on both nucleosomes, consistent with the published ubiquitin mutagenesis studies that found no other region of the ubiquitin surface to be necessary for Dot1L activation. Taken together, these results suggest that Dot1L Ile290, Leu322, and Phe326 interactions with ubiquitin are critical to ubiquitin-dependent activity. Similar to the nucleosome-interaction loop, sequences from Dot1L orthologs in higher eukaryotes align well with human Dot1L for positions at the ubiquitin interface (Figure 3C).

**Figure 3.**
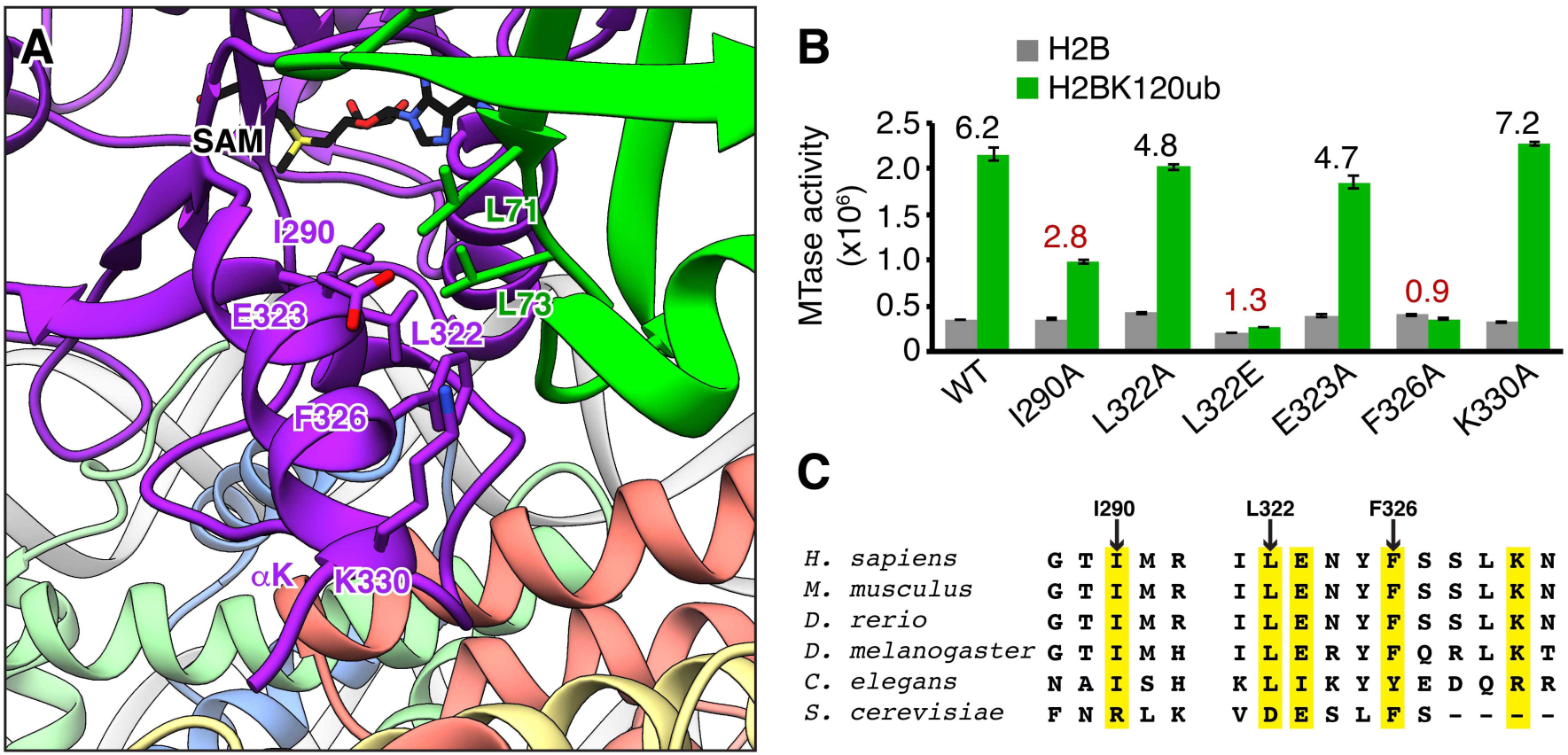
Ubiquitin interacts with Dot1L to enhance H3K79 methylation. **A**, Zoomed view of Dot1L-ubiquitin interaction with relevant side chains indicated. **B**, Quantified methyltransferase assay of Dot1 mutants on unmodified (H2B, grey) and ubiquitylated (H2BK120ub, green) nucleosomes. Numbers above bars indicate fold-enhancement by ubiquitin for each Dot1L protein. A value of 1 indicates no difference in activity toward H2BK120ub relative to unmodified nucleosomes. Five replicates performed for all assays and means and standard deviations are shown. **C**, Sequence alignment of Dot1L ubiquitin-binding regions with key positions highlighted in yellow.

### Dot1L active state model

We believe our structure represents a poised conformation of Dot1L on the nucleosome. Our biochemical analysis shows that key interactions observed in the poised state are also required for activity and are thus likely to interact during catalysis. Therefore, we modeled the active state of the Dot1L-H2BK120ub complex by fixing the position of the Dot1L Arg282 Cα and rotating Dot1L toward H3K79 on the nucleosome surface (Figure 4). As no non-SET domain methyltransferase structures exist in complex with a cofactor and a lysine-containing substrate, we used the distance between the SAH cofactor and the target lysine Cα in a ternary complex of the SET-domain containing SET8 methyltransferase to guide active state modeling (Couture et al., 2005). This SET8 structure shows an 8.8 Å distance between the Cα of the target lysine and the sulfur atom of the cofactor. A 17° rotation of Dot1L about Arg282 was required to replicate this distance in the Dot1L-nucleosome complex while placing the Cα of H3K79 at the entrance to the putative Dot1L substrate channel. This active state results in only minor steric clashes between Dot1L flexible loops and the nucleosome surface. Future structural characterization of an active complex will illuminate further molecular details.

**Figure 4.**
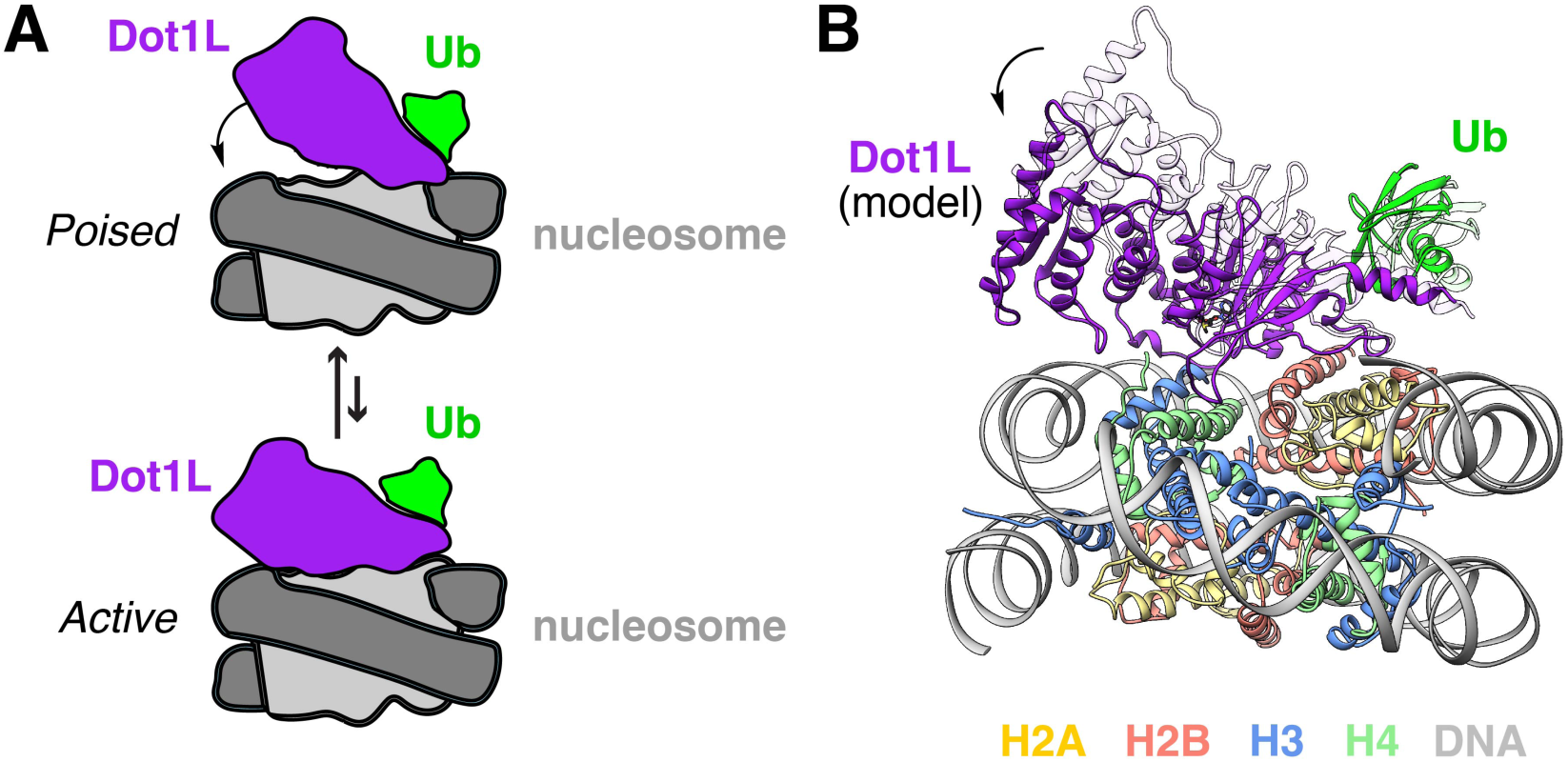
Model of active conformation of Dot1L on the nucleosome. **A**, Scheme of proposed poised to active conformational rearrangement of Dot1L and ubiquitin on the nucleosome. **B**, Manual model of Dot1L and ubiquitin in an active conformation made by a 17° rotation of Dot1L while maintaining the position of the critical Arg282 Cα atom of Dot1L. Ubiquitin was adjusted with Dot1L, keeping the orientation of ubiquitin relative to Dot1L constant. This model places a SAM cofactor in the Dot1L active site at an optimal distance to the H3K79 target. Cryo-EM model shown in white for comparison.

## Discussion

We used cryo-EM to solve the structure of Dot1L bound to an H2BK120ub nucleosome in a poised state. In our structure, Dot1L binds to the nucleosome acidic patch using a nucleosome-interaction loop. The acidic patch is emerging as a hot-spot for nucleosome binding (Kalashnikova et al., 2013; McGinty and Tan, 2016). Examination of structures of proteins and peptides bound to the nucleosome show that a common feature is the use of two arginines to bind to several locations within the acidic patch (Figure S8). The first example was observed in a crystal structure of the LANA sequence bound to the nucleosome (Barbera et al., 2006). In this structure, LANA Arg9 inserts into a cavity surrounded by H2AE61, D90, and E92 side chains. Other structures, including those with RCC1, Sir3, 53BP1, RING1B, and CENP-C, have demonstrated that many more proteins use arginines with nearly identical conformations to bind in this same acidic patch cavity, leading to the introduction of the term arginine anchor (Armache et al., 2011; Kato et al., 2013; Makde et al., 2010; McGinty et al., 2014; Wilson et al., 2016). All of these proteins also engage the nucleosome with one or more additional arginines that bind to the acidic patch in other locations. Other structures, including our Dot1L-nucleosome structure presented here and a structure of the SAGA deubiquitinating (DUB) module (Morgan et al., 2016), have nucleosome interactions with multiple arginines and no binding in the H2AE61/D90/E92 cavity. In general, the arginines that bind these other acidic patch surfaces exhibit more diversity in both binding sites and arginine side chain conformations than those that bind to the H2AE61/D90/E92 cavity. We suggest the use of the terms canonical arginine anchor and variant arginine anchor to describe these two classes of arginines. The Dot1L variant arginine anchor Arg282 most closely resembles the CENP-C variant arginine anchor Arg714. While we do not observe density for Dot1L Arg278 in our cryo-EM map, the position of its Cα atom is similar to that of the LANA variant arginine anchor Arg12. Overall, the analysis of diverse acidic patch binding proteins demonstrates both shared and protein-specific molecular mechanisms for nucleosome binding in this region.

Interestingly, we found that the LANA sequence is unable to bind to H2BK120ub nucleosomes under conditions where binding to unmodified nucleosomes was observed. This suggests that in addition to promoting Dot1L activity in a purified system, H2BK120ub may also prevent other proteins from binding the nucleosome in a physiologic cellular environment. This would limit competition, further promoting intranucleosomal methylation and may also increase local concentrations of Dot1L in areas of the nucleus containing H2BK120ub-modified chromatin. Macromolecular crystals of H2BK120ub nucleosomes do not alter the nucleosome structure and lack density for ubiquitin, suggesting ubiquitin has conformational flexibility in the absence of other bound factors (Machida et al., 2016). Therefore, it is unlikely that ubiquitin competes with LANA by binding directly to the acidic patch. Rather we favor the explanation that a conformationally flexible ubiquitin in the vicinity of the acidic patch can prevent acidic patch binding of other proteins. Some proteins for which H2BK120ub inhibits chromatin binding may have been identified in nucleosome affinity proteomics experiments with ubiquitylated nucleosomes, including RCC1, another acidic patch binder (Shema-Yaacoby et al., 2013). H2BK120ub is one of many acidic patch proximal histone modifications, and several of these modifications also tune the nucleosome binding and/or activity of chromatin effector proteins (Dann et al., 2017; Fujiki et al., 2011; Jbara et al., 2016). Given the emerging trend of acidic patch-dependent chromatin binding, this is likely to be a major mechanism in the regulation of chromatin processes.

In our structure, ubiquitin is lifted off the nucleosome surface and is in a position to interact with the catalytic domain of Dot1L. Computational sorting of particles with different conformational states of Dot1L with respect to the nucleosome demonstrates that Dot1L and ubiquitin move together through a continuum of positions. We also identify mutants in Dot1L that preferentially disrupt activity on H2BK120ub nucleosomes over unmodified nucleosomes. Taken together, our data suggest that ubiquitin enhances Dot1L activity through direct interactions with the catalytic domain of Dot1L. Contrary to this model, photocrosslinking studies place ubiquitin Leu71 and Leu73 in proximity to the H2A N-terminal tail (Zhou et al., 2016). We have not observed any density for ubiquitin near the emergence of the H2A tail from the nucleosome disk or for the first 15 residues of the H2A tail either interacting with ubiquitin or other parts of the complex. As our cryo-EM map predicts conformational flexibility in ubiquitin, we anticipate that crosslinking to H2A may occur only in minor ubiquitin conformations. Previous results show that moving ubiquitin from H2BK120 to H2BK125 leads to a modest decrease in Dot1L activity, while moving ubiquitin to H2BK116 or K108 dramatically or entirely inhibits Dot1L, respectively (Supplementa1 Figure S9) (Chatterjee et al., 2010). While we do not see clear density for the ubiquitin-H2BK120 linkage in our cryo-EM map, we hypothesize that the ubiquitin C-terminal tail is flexible enough to allow ubiquitin to reach its observed position in our structure from an H2BK125 attachment site. However, H2BK108 and H2BK116 attachment places ubiquitin between Dot1L and the nucleosome and likely prevents productive binding of Dot1L as a result.

We generated a model of Dot1L in an active conformation on the nucleosome surface. This model largely accommodates Dot1L loops without structural rearrangements. It has been demonstrated that a basic sequence in the H4 N-terminal tail is critical for Dot1L activity (Fingerman et al., 2007; McGinty et al., 2009). In our model, there is an acidic cleft on the surface of Dot1L adjacent to the location from which the H4 N-terminal tail emerges from the nucleosome. We hypothesize that the H4 basic patch binds to this Dot1L acidic cleft to promote formation of a catalytically active conformation of Dot1L on the nucleosome.

In summary, we solved the cryo-EM structure of Dot1L bound to an H2BK120ub nucleosome. Dot1L binds to the nucleosome acidic patch using a nucleosome-interaction loop and interacts directly with ubiquitin for activation. Dot1L-mediated methylation of H3K79 is essential to the pathogenesis of MLL-rearranged leukemias, prompting the development of Dot1L inhibitors that are now in clinical trials. Our structure and subsequent biochemical validation identify mutations to interrogate the nucleosome-specific and ubiquitin-dependent functions of Dot1L. This will enable precision hypothesis-driven analysis of the functions of Dot1L in normal and leukemia model cells. Further, our studies suggest that H2BK120 ubiquitylation may function as a gatekeeper of the acidic patch. The tuning of acidic patch binding by adjacent post-translational modifications is likely to be a fundamental mechanism in the regulation of chromatin signaling.

## Materials and Methods

### Preparation of proteins and nucleosomes

An unpublished plasmid with the gene fragment encoding human Dot1L(2–416) (Dot1L_cat_) in the pST50Tr vector with an N-terminal Strep peptide (STR)-hexahistidine (His_6_) affinity tag was obtained as a gift from Song Tan. STR-His_6_-hDot1L_cat_ was expressed in *E. coli* BL21(DE3)pLysS cells at 18°C. The tagged protein was enriched by metal-affinity chromatography using Talon resin (Clontech). The affinity tag was removed using the tobacco etch virus (TEV) protease prior to further purification by cation exchange chromatography using a Source S resin (GE Healthcare). For structural biology, an additional gel filtration chromatography step with a Superdex 200 increase 10/300 column (GE Healthcare) was performed. All Dot1L_cat_ point mutants were cloned by site-directed mutagenesis and expressed and purified identically to the wild-type protein fragment. All Dot1L mutants were verified by intact mass spectrometry using an Agilent 6520 Accurate Mass Quadrupole Time-of-Flight mass spectrometer (Figure S10). Unpublished pST50Tr plasmids containing the sequence encoding LANA peptide residues 1–23 or a triple mutant of residues 8–10 (LRS to AAA) with an N-terminal AviTag (AVT)-glutathione-S-transferase (GST)-His_6_ tag were obtained as a gift from Jiehuan Huang and Song Tan. AVT-GST-His6-LANA and -LANA(LRS>AAA) were expressed in *E. coli* BL21(DE3)pLysS cells at 37°C. The fusion proteins were purified by Talon metal-affinity chromatography and anion exchange chromatography using a Source Q resin (GE Healthcare). An unpublished pST50Tr plasmid containing STR-His_10_-tagged *E. coli* exodeoxyribonucleaseX (ExoX) residues 2–167 with an Asp6Ala inactivating mutant was a gift from Song Tan. This inactive ExoX was expressed and purified as described above for Dot1L_cat_. TEV cleavage leaves a 3x Gly-Ser sequence at the N-terminus of ExoX.

Recombinant *Xenopus* and human histones were expressed, purified and reconstituted into nucleosomes as previously described (Luger et al., 1999). Nucleosomes were assembled with 147 or 155 bp of the 601 DNA sequence (Lowary and Widom, 1998) with the nucleosome positioning sequence centered in each DNA fragment. H2BK120ub(DCA) was prepared essentially as previously described (Morgan et al., 2016). Briefly, *Xenopus* H2BK120C was cloned by site-directed mutagenesis and the pST50Tr-STR-His_10_-ubiquitin(Gly76Cys) plasmid was obtained as a gift from Ryan Henrici and Song Tan. Purified *Xenopus* H2A/H2BK120C dimer and purified STR-His_10_-ubiquitin(Gly76Cys) were combined at 0.5 and 1 μM concentrations, respectively, in crosslinking buffer (6 M urea, 33.3 mM sodium borate pH 8.5, 5 mM TCEP) and cooled on ice. Crosslinking was performed by the addition of dichloroacetone (DCA) to a final concentration of 2.5 μM. After 60 min on ice, DCA was quenched with the addition of 50 mM β-mercaptoethanol. Histone dimer was refolded from the mixture as previously reported and purified with Talon resin to remove any H2A/H2BK120C lacking DCA-crosslinked ubiquitin. TEV protease was used to remove the STR-His_10_ tag on ubiquitin, and H2BK120ub(DCA)-containing histone dimer was purified away from crosslinked ubiquitin dimers by cation exchange chromatography. A Gly-Ser-Gly-Ser sequence remains on the N-terminus of ubiquitin after protease cleavage. *Xenopus* H2BK120ub(Gly76Ala) was prepared as previously described (McGinty et al., 2009). Mass spectrometric characterization was preformed using an Agilent 6520 Accurate Mass Quadrupole Time-of-Flight mass spectrometer. H2BK120ub analogs were assembled into nucleosomes as described above.

### Reconstitution of Dot1L_cat_-H2BK120ub nucleosome complex

The Dot1L_cat_-H2BK120ub(DCA) nucleosome complex used for cryo-EM experiments was reconstituted as previously reported for other nucleosome complexes (McGinty et al., 2016). Briefly, 2.5 molar equivalents of Dot1L_cat_ were added in four steps to H2BK120ub(DCA) nucleosomes containing all *Xenopus* histones and 147 bp 601 DNA at 5 minute intervals in reconstitution buffer (25 mM Tris-Cl pH 7.6, 150 mM NaCl, 1 mM DTT, 0.1 mM PMSF). The complex was purified by gel filtration using a Superdex 200 increase column equilibrated in reconstitution buffer. For crystallography experiments, Dot1L_cat_ was reconstituted using DCA-crosslinked H2BK120ub nucleosomes containing 155 bp 601 DNA and the inactive ExoX as a crystallization chaperone. Reconstitution of the complex was performed as described above, with the following changes: 1) 2.5 molar equivalents of ExoX was added in four steps after addition of Dot1L and 2) reconstitution and purification were performed in reconstitution buffer containing 50 mM NaCl.

### Cryo-EM sample preparation

Dot1L_cat_-H2BK120ub(DCA) nucleosome complex at ~10 mg/ml was buffer exchanged into crosslinking buffer (10 mM HEPES pH 7.5, 150 mM NaCl) using a Zeba spin desalting column (Thermo Fisher). Glutaraldehyde (Sigma Aldrich) was added to a final concentration of 0.1% and crosslinking was allowed to proceed for 5 min at room temperature. Glutaraldehyde was quenched by the addition of Tris-Cl at pH 7.5 to a final concentration of 20 mM. After 15 min at room temperature, samples were buffer exchanged into crosslinking buffer. Complex was diluted to 0.93 mg/ml in freezing buffer (10 mM HEPES pH 7.5, 48 mM NaCl) prior to spotting on a Quantifoil R1.2/1.3, 300 mesh grid for vitrification after blotting for 4 s using a Vitrobot Mark IV (FEI) cryoplunger at 100% humidity and 4°C. An identical complex was analyzed by liquid chromatography coupled mass spectrometry using an Agilent 6520 Accurate Mass Quadrupole Time-of-Flight mass spectrometer to assess the methylation state of histone H3.

### Cryo-EM data collection and analysis

Cryo-EM data was collected on a Titan Krios (FEI) operating at 300 kV. The images were recorded using the automated data acquisition software EPU (FEI) using a Falcon III direct electron detector (FEI) operated in electron counting mode at a nominal magnification of 75,000, corresponding to a pixel size of 1.08 Å, and at a defocus range of −1.25 μm to −2.75 μm. Each exposure lasted 60 s and was collected as a 30-frame movie at a dose rate of ~0.8e^−^/pix/s, resulting in a total electron dose of ~42 e^−^/Å^2^. Whole-frame movie alignment was performed with MotionCor2 (Zheng et al., 2017), CTF parameters were estimated with GCTF (Zhang, 2016), and Relion v2.1 (Kimanius et al., 2016) was used for subsequent data analysis. Initially, ~1,000 particles were manually picked to generate 2D class averages to use as templates for iterative rounds of automatic particle picking. In total, 408,526 automatically selected particles were subjected to reference-free 2D classification. Iterative rounds of 2D classification were used to narrow the set to 156,709 high quality particles. 3D classification using a 50 Å low pass filtered nucleosome structure (PDBID 6ESF) generated in Relion as reference identified two classes (3 and 4) containing clearly defined Dot1L-nucleosome complexes. Movie refinement, particle polishing, 3D refinement, and post-processing of classes 3 and 4 combined resulted in a 3.5 Å reconstruction showing significant conformational heterogeneity in Dot1L and ubiquitin. Polished particles from classes 3 and 4 were subjected to further 3D classification into 4 new subclasses (designated with A). Refinement and post-processing of the resultant class 2A yielded a 3.9 Å reconstruction used for modeling. Masking of the Dot1L and ubiquitin densities for class 2A and continuing refinement resulted in a 7.6 Å map with improved density for Dot1L and ubiquitin. Finally, the original class 3 and class 4 polished particles were reclassified into 10 3D subclasses (designated with B), allowing further parsing of stoichiometry and conformational heterogeneity. Crystal structures for the nucleosome (PDBID 3LZ0), the catalytic domain of Dot1L (PDBID 1NW3), and ubiquitin (PDBID 1UBQ) were docked into the 3.9 Å reconstructed map using Chimera (Pettersen et al., 2004) and refined by iterative real-space refinement in Phenix (Afonine et al., 2018) and manual model building in Coot (Emsley et al., 2010). Local resolution was calculated with ResMap (Kucukelbir et al., 2014). Resolution of reconstructions was calculated using the gold standard FSC criterion of 0.143. All molecular graphics were prepared with Chimera.

### Crystallization and X-ray crystallography

The Dot1L_cat_-ExoX-H2BK120ub(DCA) nucleosome complex was concentrated to 10 mg/ml and crystallized by mixing 1:1 with 25 mM Bis-Tris pH 6 (Hampton), 25 mM (NH_4_)_2_HPO_4_ (Hampton), and 1.5% PEG2000-MME (Fluka) at room temperature using modified microbatch under oil with Al’s oil (D’Arcy et al., 2004). Notably, this complex was not crosslinked with glutaraldehyde. Inactive ExoX was used as a crystallization chaperone (Robert McGinty and Song Tan unpublished results). Crystals were collected after 20–35 days and dehydrated and cryoprotected by soaking at room temperature in 25 mM Bis-Tris pH 6, 25 mM (NH_4_)_2_HPO_4_, and 3.75% PEG2000-MME in drops containing increasing concentrations of PEG400 (final concentration 24%, increment 4%, soak interval 15 min). Upon completion of soaking, crystals were flash cooled in liquid nitrogen. Diffraction data was collected on the Advanced Photon Source SER-CAT 22-ID beamline at 1 Å and 100 K. Data was processed using HKL2000 (Otwinowski and Minor, 1997) in the C2221 space group. A molecular replacement model was generated with CCP4 Phaser MR (McCoy et al., 2007) using a polyalanine half nucleosome model generated from PDBID 1KX5 and a polyalanine Dot1L catalytic domain model generated from PDBID 1NW3, with loop residues 296–313 deleted.

### Lysine methyltransferase assays

Lysine methyltransferase assays were performed using the MTase-Glo enzyme-methyltransferase kit (Promega) using the manufacturer’s protocol. Briefly, 1 μM nucleosomes were combined with 15 nM wild-type or mutant Dot1L_cat_ in methyltransferase buffer (20 mM Tris-Cl pH 8.0, 50 mM NaCl, 1 mM EDTA, 0.1 mg/ml BSA, 1 mM DTT, and 20 μM SAM) in a total volume of 8 μl. The concentration of 15 nM Dot1L was selected as it was very close to the linear range of activity on H2BK120ub nucleosomes, while allowing activity on unmodified nucleosomes to be measured robustly. Using the same concentration allowed the direct comparison of activity on ubiquitylated and unmodified nucleosomes. The assay was allowed to proceed at 30°C for 30 min prior to developing the assay and quantifying luminescence using an Envision 2103 Multilabel plate reader (Perkin Elmer). Assays were performed with 5 replicates. All assays were performed with human histones except for *Xenopus* unmodified or ubiquitylated H2B. No significant difference in Dot1L activity was observed between nucleosomes containing *Xenopus* H2B and those containing canonical human H2B, validating the use of chimeric nucleosomes. For LANA competition methyltransferase activity assays, AVT-GST-His_6_-LANA or AVT-GST-His_6_-LANA(LRS>AAA) were added to the assay at 0, 5, 10, 25, 50, and 100 μM final concentrations. An additional assay was performed after incubation of LANA fusion proteins with 0.025 molar equivalents of TEV protease overnight at room temperature. Cleavage of LANA affinity tags was verified by SDS-PAGE.

### Gel filtration-based binding assay

AVT-GST-His_6_-LANA was reconstituted with H2BK120ub(Gly76Ala) or unmodified nucleosomes as described above for Dot1L-nucleosome reconstitutions at a 4:1 molar ratio (~4.5 μM nucleosome) in 10 mM Tris-Cl pH 8.0, 50 mM NaCl, and 1 mM DTT. Interaction was evaluated by separation on a Superdex 200 increase gel filtration column equilibrated in the same buffer. Chromatography fractions were analyzed by SDS-PAGE.

## Acknowledgements

We thank Andrew Thieme and Erin Blanding for contributions to reagent preparation. Brittany Ford and Fabiola Jaramillo assisted us with cryo-EM sample preparation and screening. Cryo-EM data was collected at the Shared Materials and Instrumentation Facility at Duke University, within the collaborative framework of the Molecular Microscopy Consortium. Leonard Collins at the UNC Biomarker Mass Spectrometry Facility collected mass spectrometry data. We thank the McGinty lab, Brian Strahl, and Song Tan for project discussions and comments on the manuscript. This work was supported by Searle Scholars Program and the Pew-Stewart Scholars in Cancer Research awards to R.K.M. and NIH T32GM008570 to C.J.A.

## Author Contributions

C.J.A. and R.K.M. planned experiments, collected and analyzed data, and prepared the manuscript. M.R.B. and M.J.B. planned experiments, collected and analyzed data, and edited the manuscript. A.H., Y.K., and E.K.H. collected and analyzed data. All authors have commented on and agreed to the content of this manuscript.

---

**Table S1.**
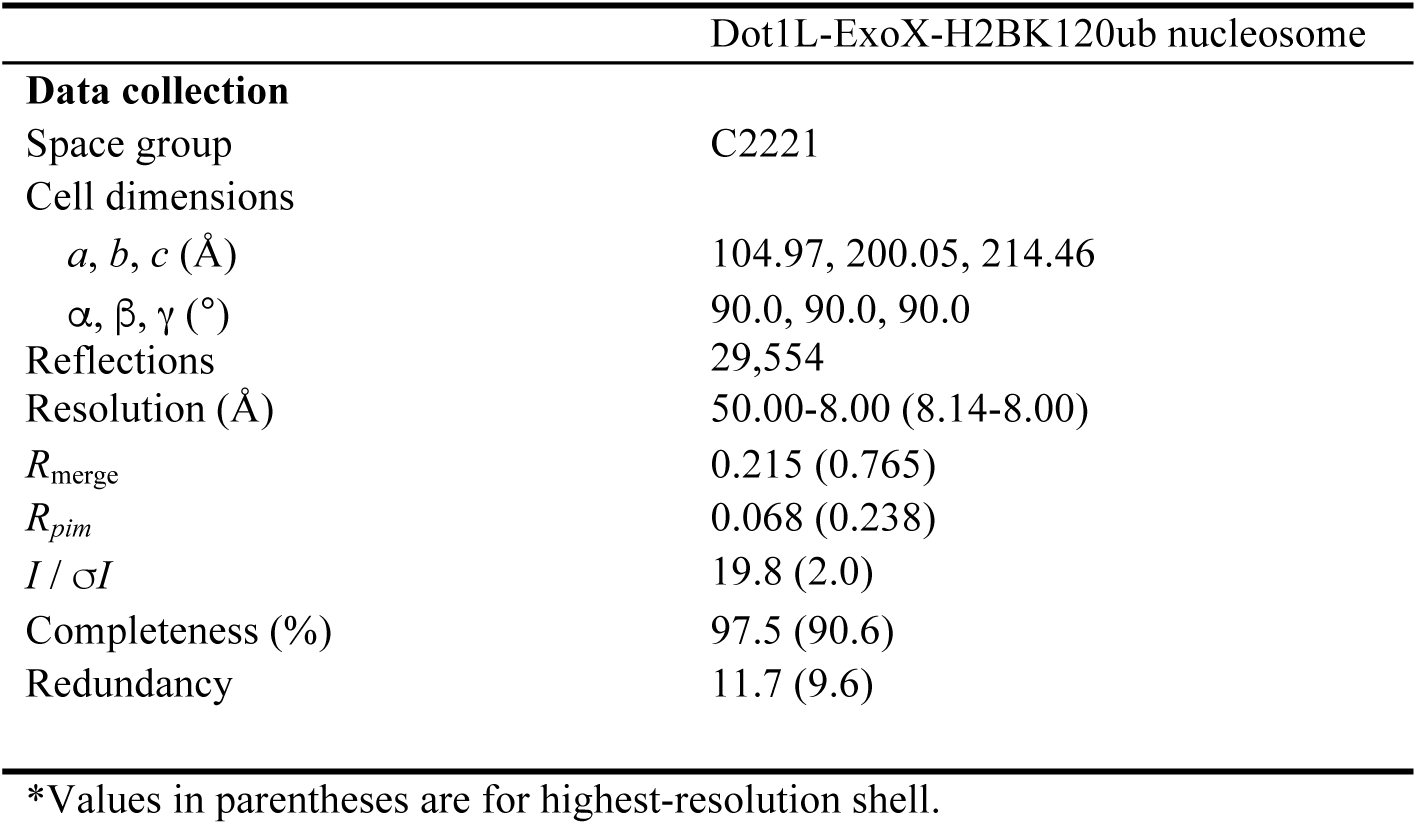
X-ray crystallography data collection

**Figure S1, related to Figure 1.**
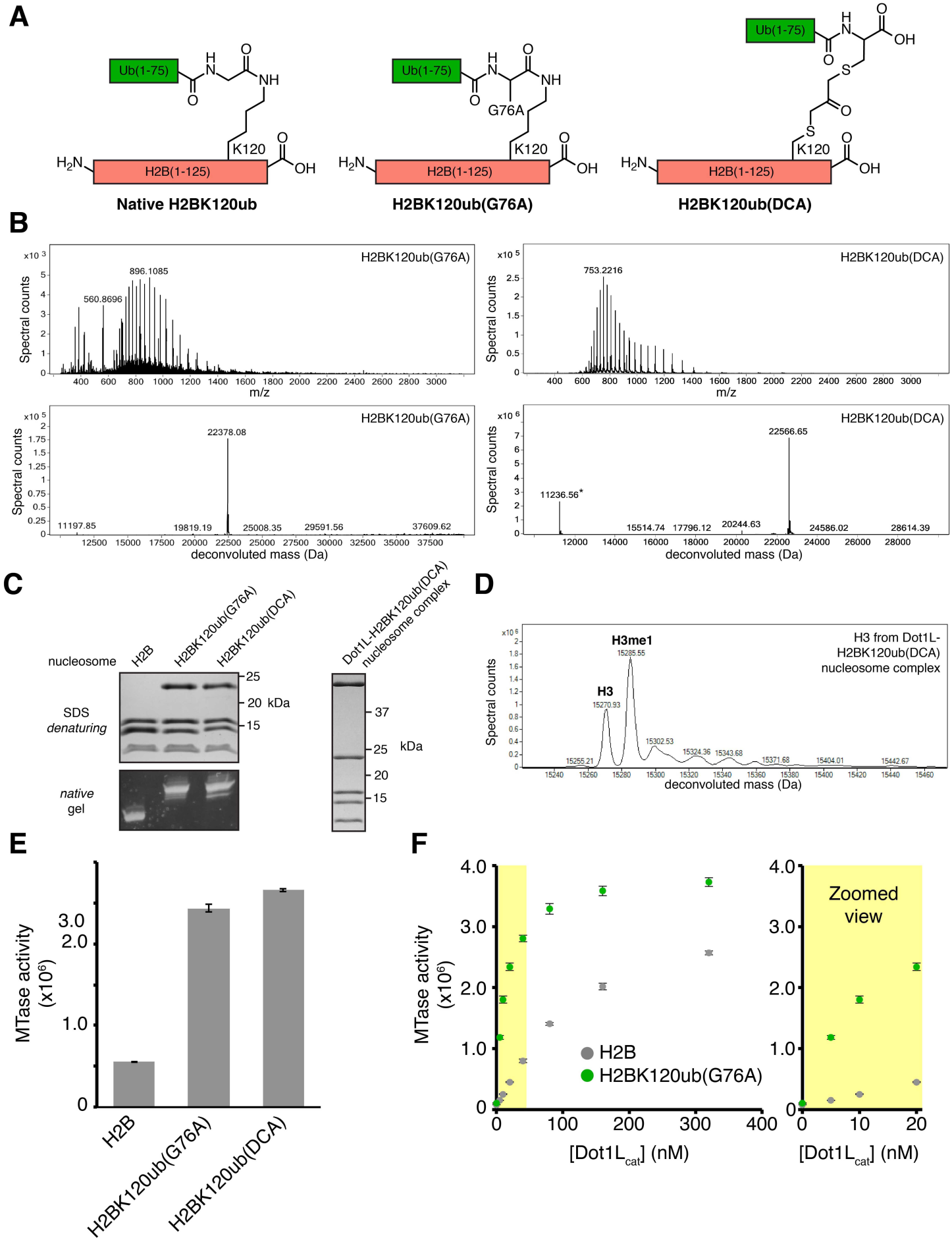
Preparation and characterization of H2BK120ub analogs. **A**, Scheme showing differences between native, G76A, and DCA-crosslinked linkages at the H2B-ubiquitin junction. **B**, Left, ESI-mass spectrum of semisynthetic H2BK120(Gly76Ala = G76A) (top) and deconvoluted mass spectrum (bottom, expected mass 22,378 Da); Right, equivalent spectra for H2BK120(DCA) from Dot1L-H2BK120(DCA) nucleosome complex (expected mass 22,566 Da; asterisk marks mass of coeluting histone H4). **C**, SDS denaturing and 10% native acrylamide gels of nucleosomes containing unmodified or ubiquitylated H2BK120 (G76A or DCA crosslinked) (left) and of reconstituted Dot1L-H2BK120ub(DCA) complex used for cryo-EM (right). **D**, Deconvoluted ESI-mass spectrum of histone H3 from Dot1L-H2BK120ub(DCA) nucleosome complex (expected mass unmethylated 15,271 Da, monomethylated 15,285 Da). **E**, Quantified methyltransferase assay using nucleosomes reconstituted with indicated unmodified or ubiquitylated H2B. **F**, Quantified methyltransferase assay with designated Dot1L_cat_ concentrations. Zoomed view of indicated region (yellow) at right. Five replicates performed for all assays and means and standard deviations are shown.

**Figure S2, related to Figure 1.**
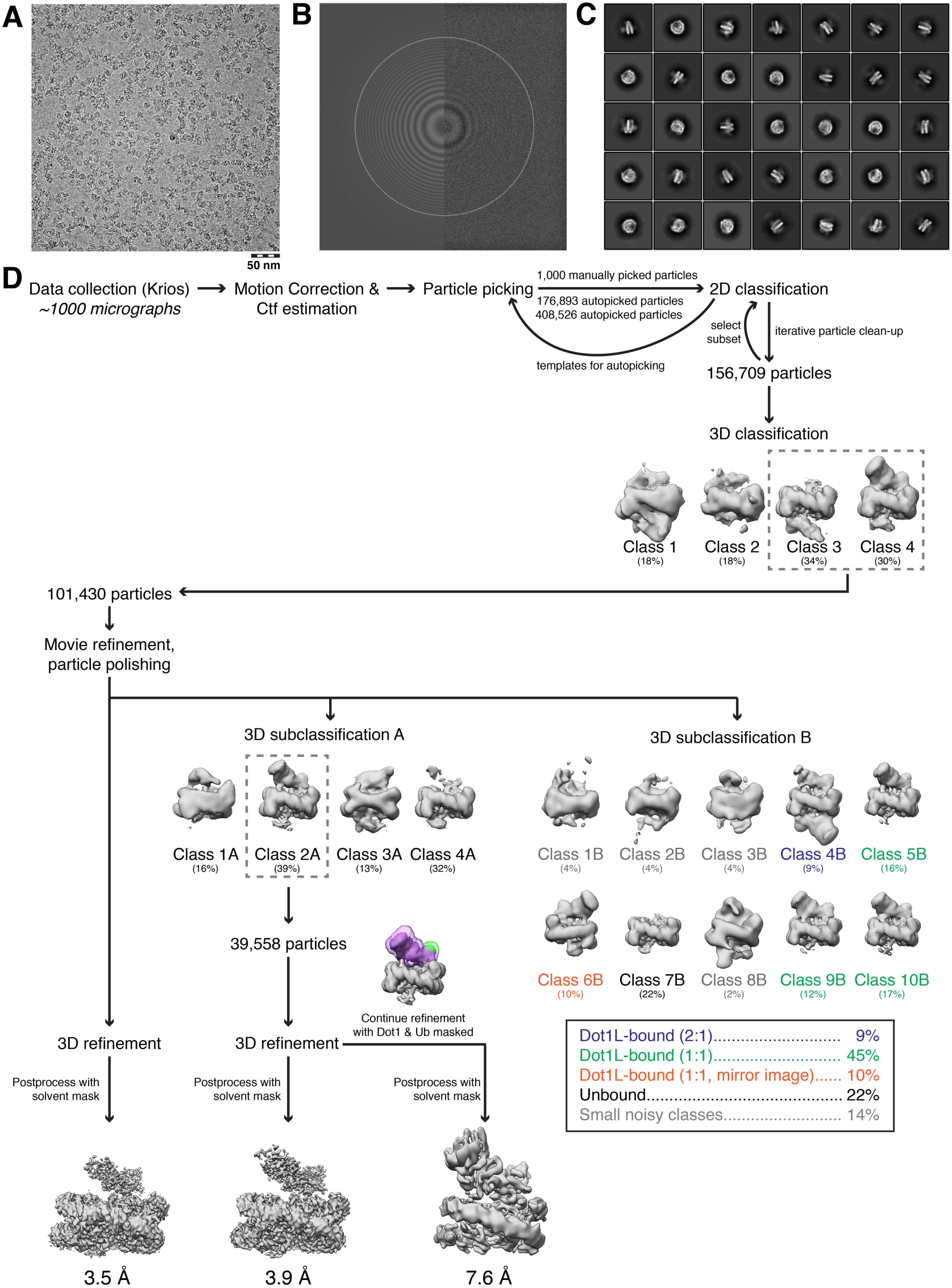
Workflow for cryo-EM data processing. Representative **A**, micrograph of complex, **B**, CTF estimation by GCTF, and **C**, reference-free 2D classifications. **D**, Data analysis scheme showing multiple classification strategies and reconstructed maps discussed in the main text. Classes 3 and 4 of initial 3D classification gave rise to a 3.5 Å map with poorly resolved density for Dot1L and ubiquitin (left). Dot1L and ubiquitin densities were improved by further classification into 4 subclasses leading to a 3.9 Å map from class 2A (middle). The Dot1L and ubiquitin volumes were masked to improve these regions of the map to allow for high confidence docking of high resolution crystal structures into the map. Finally, finer classification of the particles from the initial classes 3 and 4 into 10 subclasses was used to parse stoichiometric and conformational heterogeneity (right).

**Figure S3, related to Figure 1.**
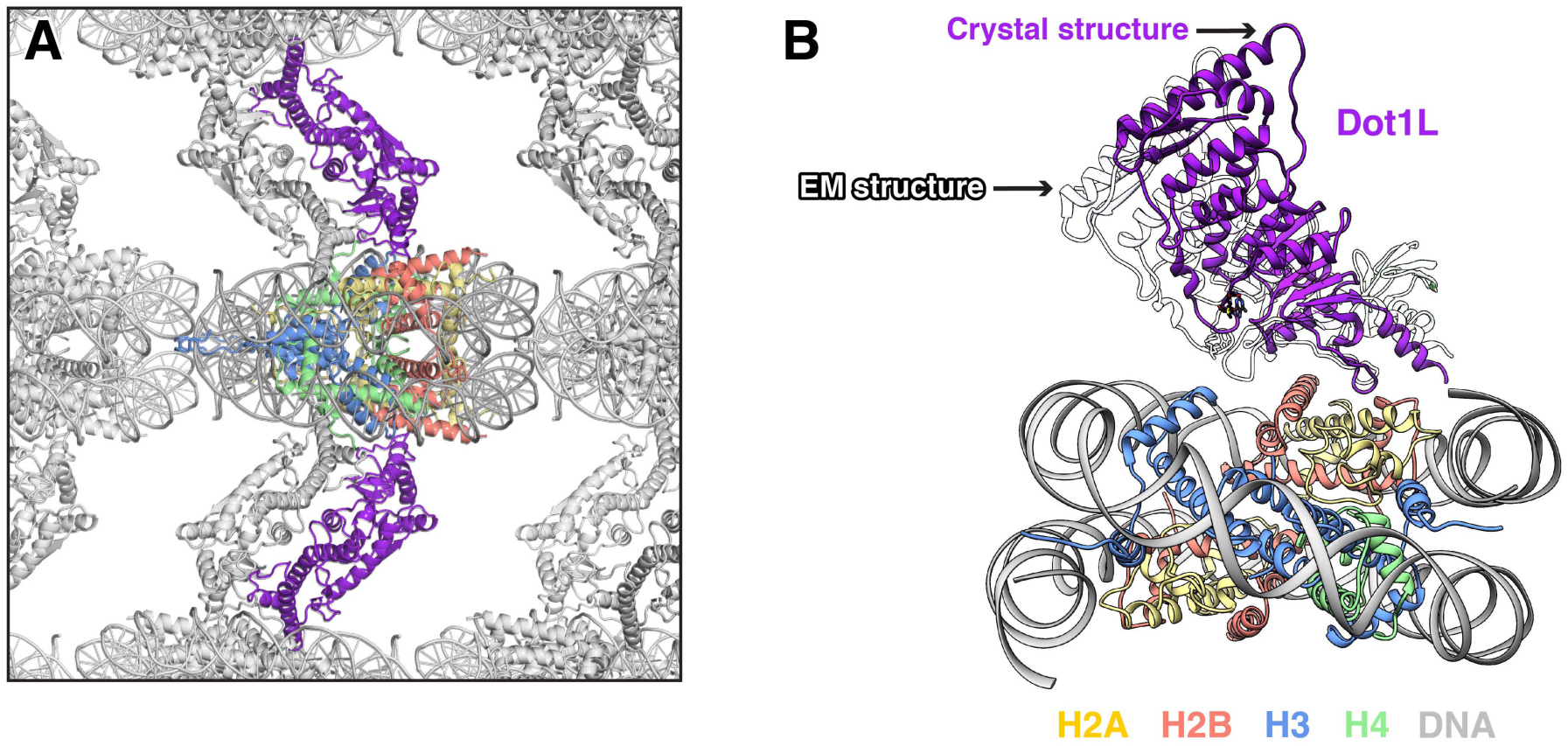
Comparison of X-ray and cryo-EM models of Dot1L-nucleosome complex. **A**, Symmetry-related complexes in crystal used for molecular replacement model with one nucleosome and two Dot1Ls colored. Another Dot1L from an adjacent unit cell wedges between Dot1L and the nucleosome to which it is bound on each nucleosome face. **B**, Alignment of cryo-EM and X-ray molecular replacement models using histones for alignment of structures. Dot1L from crystal model (purple) is lifted slightly away from the nucleosome as compared with Dot1L from the cryo-EM structure (white). The nucleosome is only shown for the cryo-EM structure for simplicity.

**Figure S4, related to Figure 1.**
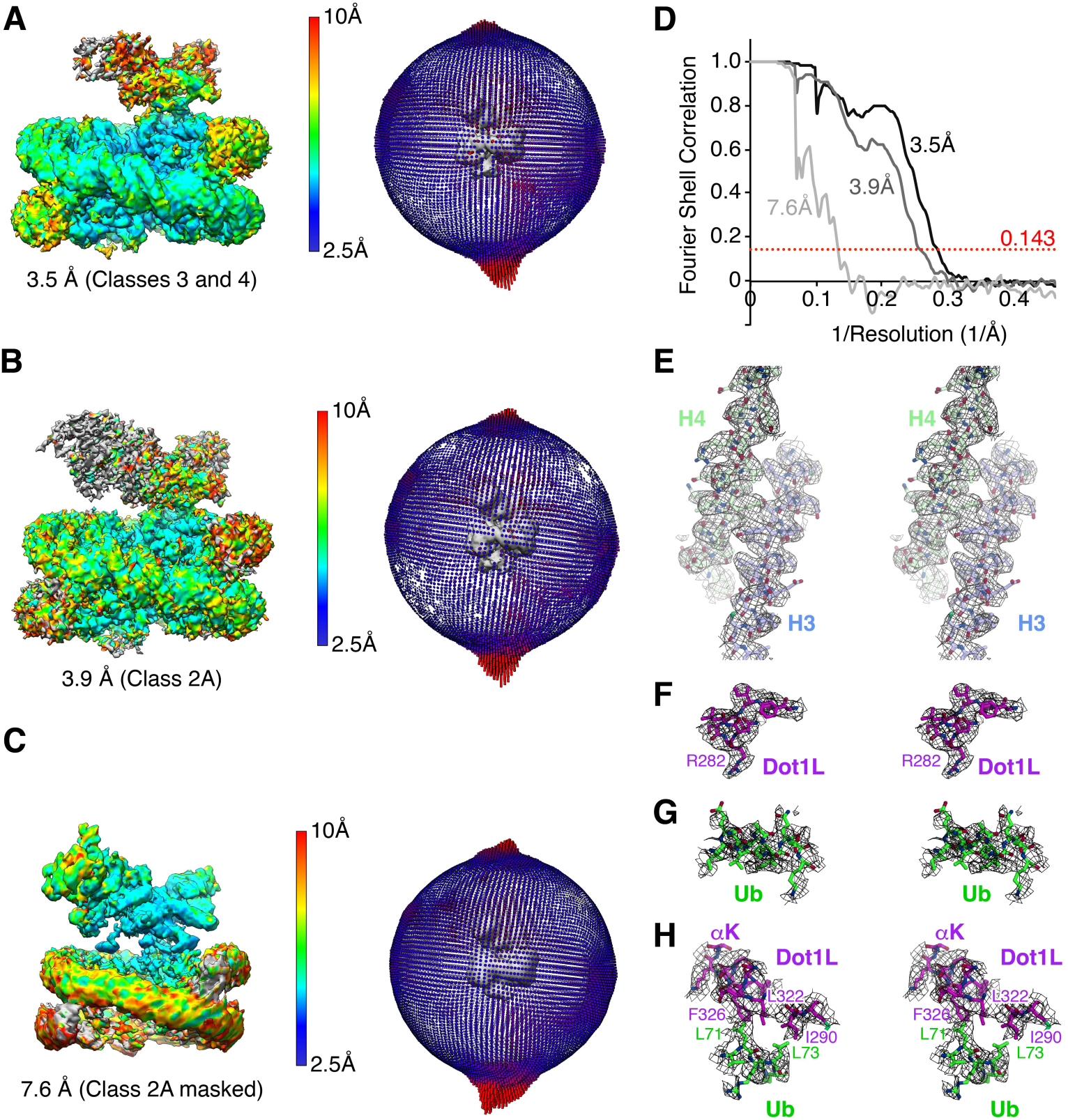
Local and overall resolution of reconstructed maps. **A**, Reconstructed map (3.5 Å) colored with local resolution (left), and orientational views (right). **B** and **C**, As in A for 3.9 Å reconstructed map and 7.6 Å reconstructed map after masking of Dot1L and ubiquitin volumes. **D**, FSC curves for post-processed reconstructions. **E-H**, Stereo views of local EM density maps from 3.9 Å reconstruction of **E**, H3 residues 89–112 and H4 residues 51–76, **F**, the Dot1L nucleosome interaction loop residues 277–284, **G**, ubiquitin residues 23–33, and **H**, Dot1L-ubiquitin interface including Dot1L residues 289–291+322–331 and ubiquitin residues 70–74. Masked .mrc maps in E-H contoured at 8 with 2.5 Å carve in PyMOL.

**Figure S5, related to Figure 1.**
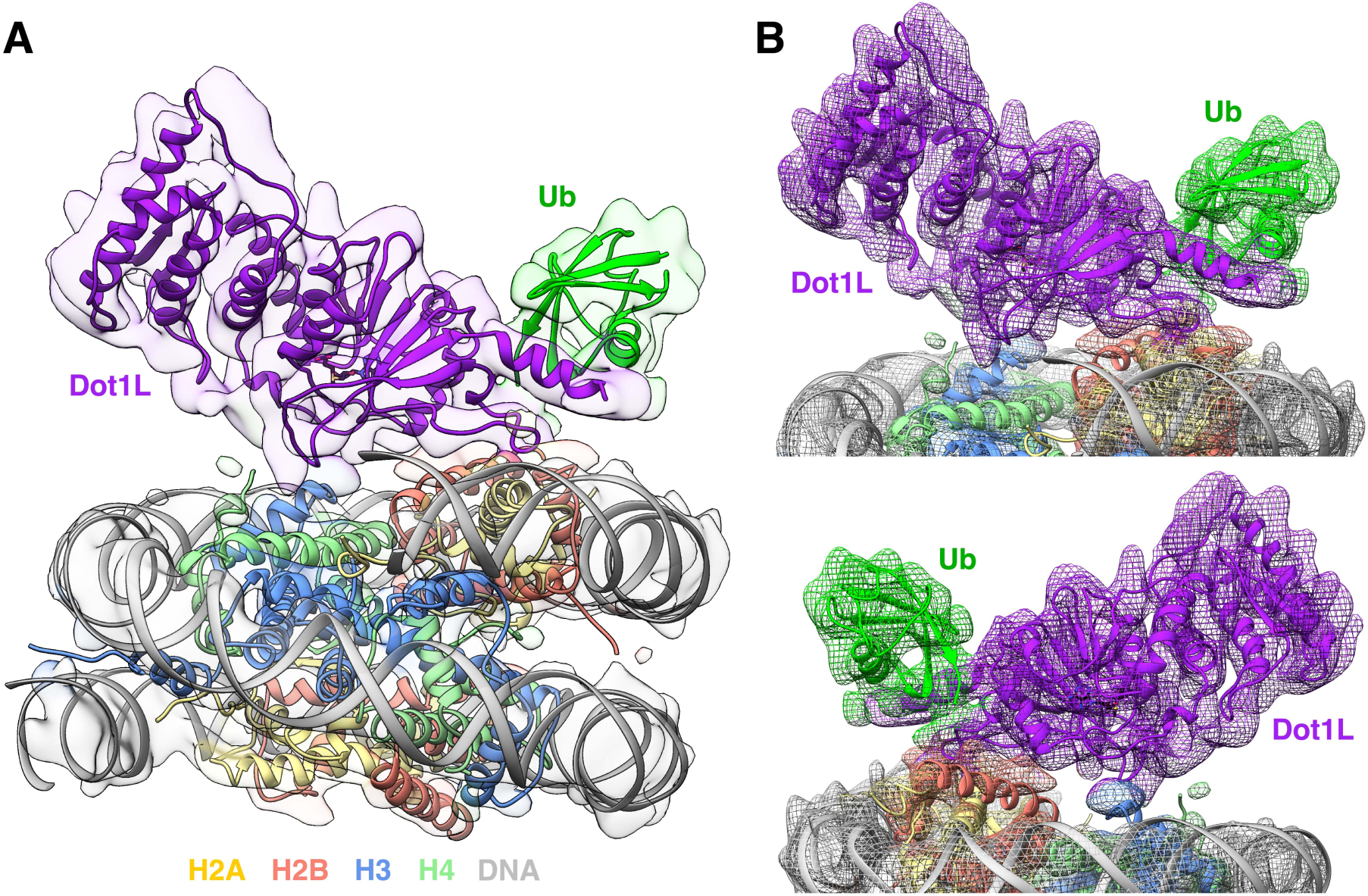
Masking of Dot1L and ubiquitin volumes confirms placement in model. **A**, Final model overlaid with 7.6 Å reconstruction from masking of Dot1L and ubiquitin volumes in Class 2A. **B** and **C**, Front and back zoomed views of overlay in panel A.

**Figure S6, related to Figure 2.**
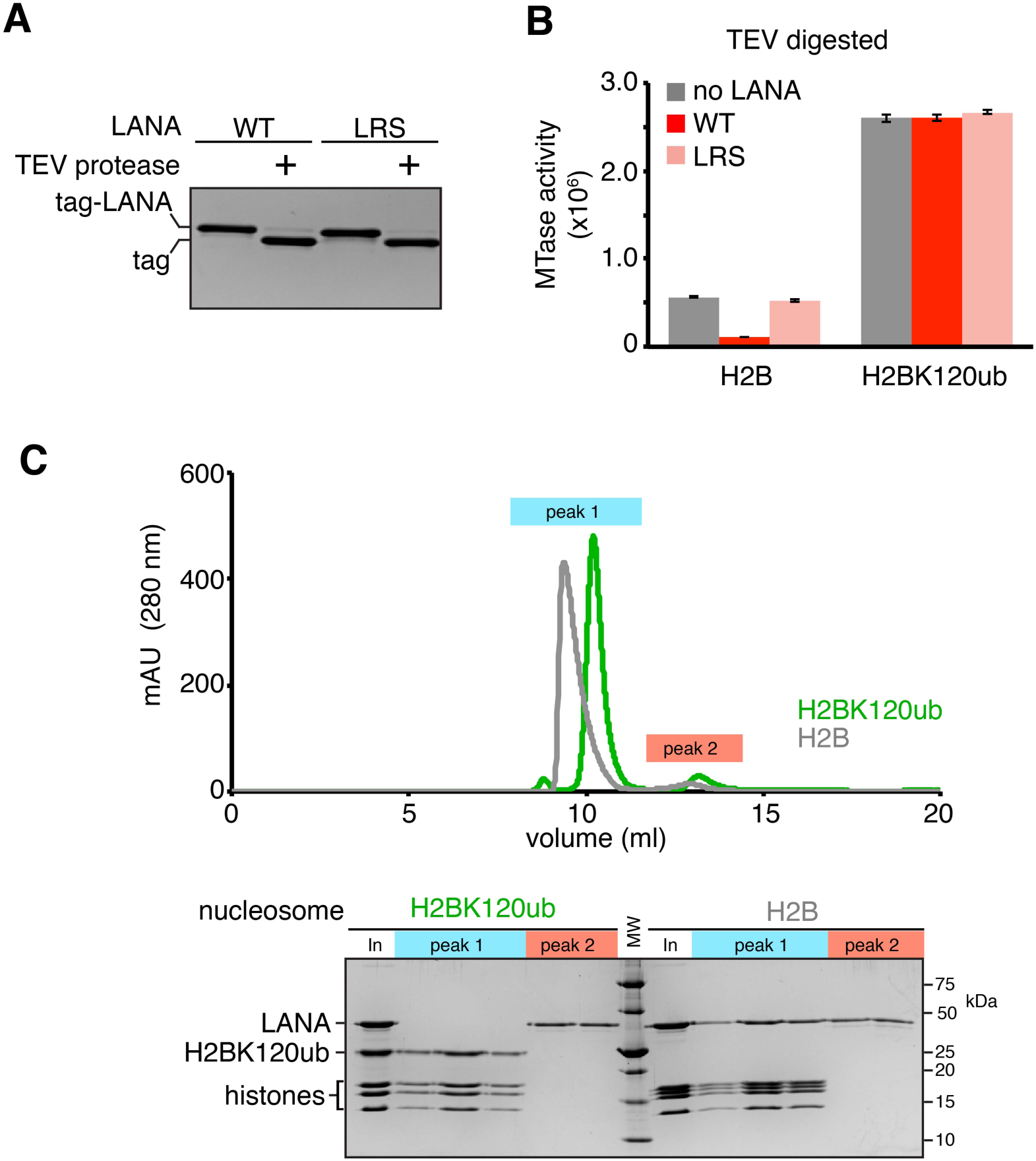
LANA cannot bind H2BK120ub nucleosomes. **A**, Denaturing gel showing TEV cleavage of LANA fusions prior to assay in B. Cleaved LANA is not observed on gel due to its small size. **B**, Quantified methyltransferase assay using no LANA or TEV cleaved wild-type or nucleosome binding-deficient (LRS = LRS 8–10 AAA) LANA fusion proteins on unmodified and H2BK120ub nucleosomes. **C**, Overlaid gel filtration chromatograms of reconstituted LANA-unmodified nucleosome complex and failed LANA-H2BK120ub nucleosome reconstitution (top). Peak 1 contains nucleosome or nucleosome-LANA complex and peak 2 contains free LANA fusion protein. Gel of representative fractions from each peak is shown (bottom). Five replicates performed for all assays and means and standard deviations are shown. In = Input.

**Figure S7, related to Figure 3.**
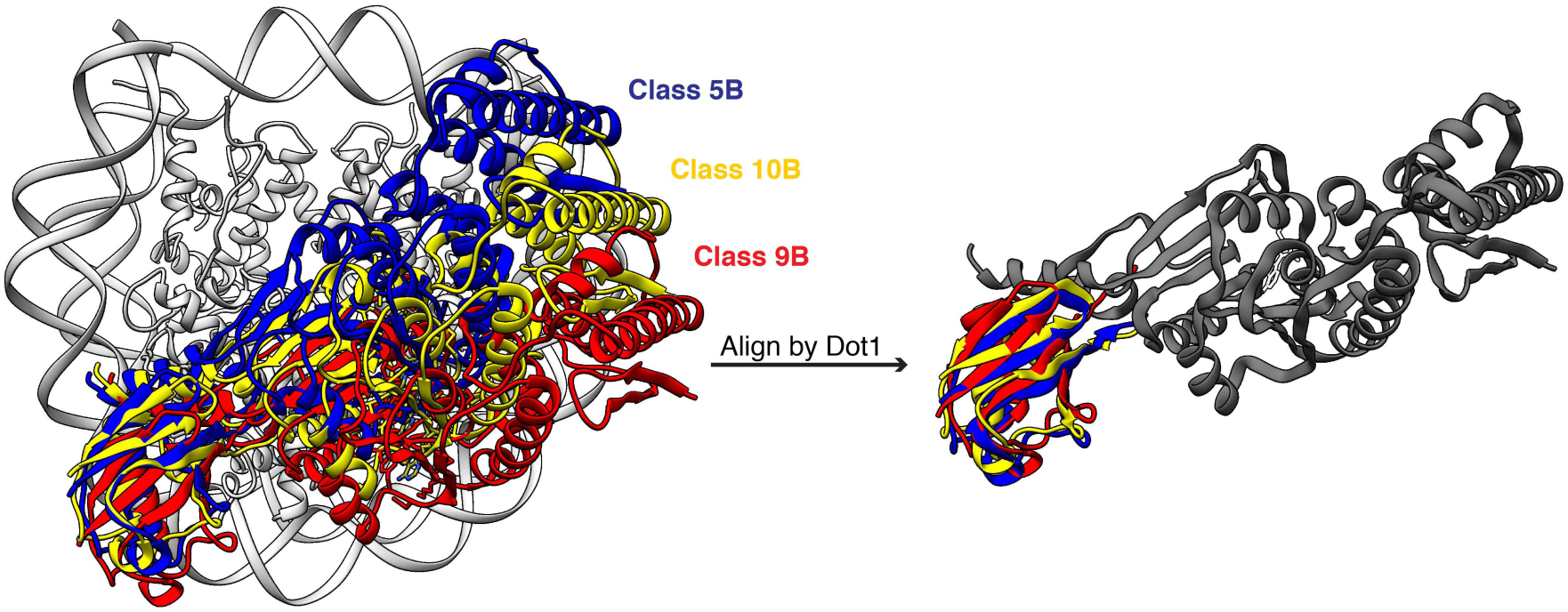
Dot1L and ubiquitin conformational heterogeneity is interdependent. Overlay of docked structures for subclasses 5B, 9B, and 10B with Dot1L and ubiquitin colored (left). Alignment of Dot1L in these classes show that the ubiquitin maintains a similar orientation relative to Dot1L (right).

**Figure S8.**
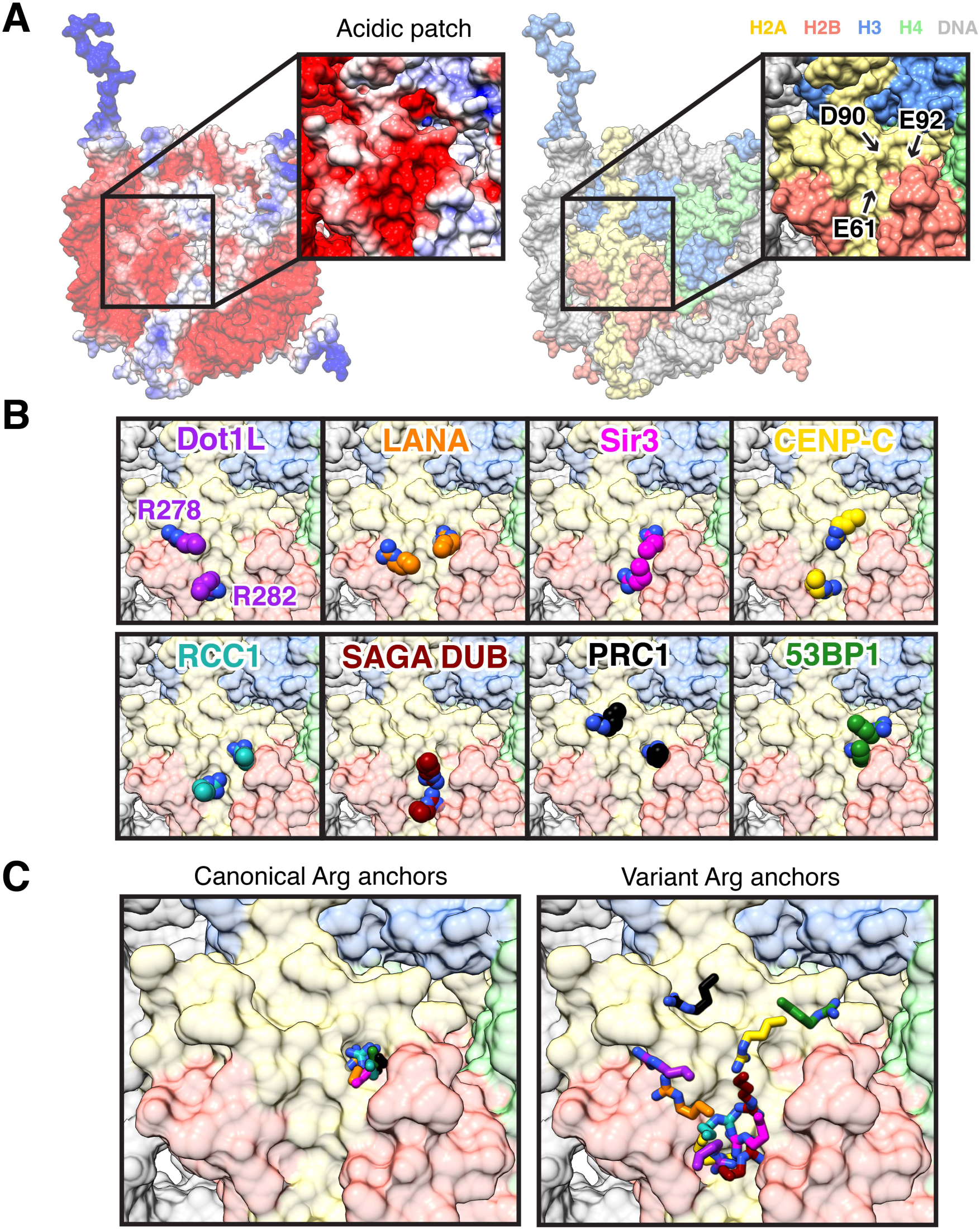
Canonical and variant arginine anchors bind the nucleosome acidic patch. **A**, Electrostatic (generated with APBS using AMBER) and histone-colored surfaces of nucleosome. The locations of H2A E61, D90, and E92 are shown. **B**, Zoomed views of arginine anchors of Dot1L, LANA (PDBID 1ZLA), Sir3 (PDBID 3TU4), CENP-C (PDBID 4X23), RCC1 (PBDID 3MVD), PRC1/RING1B (PDBID 4RP8), SAGA DUB module (PDBID 4ZUX), and 53BP1 (PDBID 5KGF). **C**, Overlay of all canonical (left) and variant (right) arginine anchors. Canonical arginine anchors bind with near identical conformation while variant arginine anchors bind with more conformational and positional diversity.

**Figure S9.**
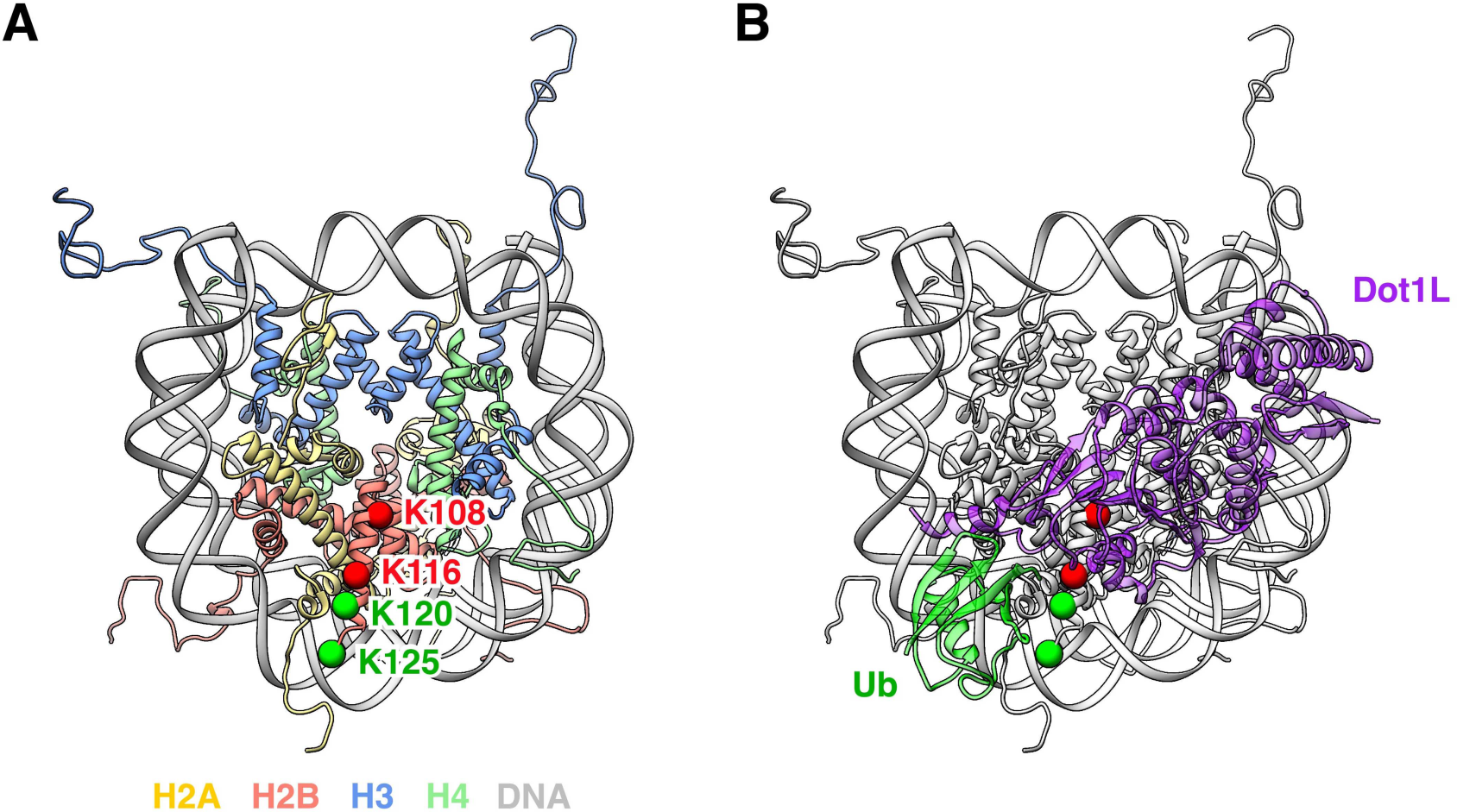
Importance of ubiquitylation position for Dot1L activity. **A**, Top view of nucleosome (PDBID 1KX5) with positions of Cα atoms from H2BK108, K116, K120, and K125 indicated. Dot1L is activated by ubiquitin attachment at green but not red sites. 1KX5 was used because H2BK125 is not observed in Dot1L-bound structure. **B**, Same view with Dot1L and ubiquitin from cryo-EM structure overlaid and displayed with transparency to allow visualization of underlying histone positions.

**Figure S10.**
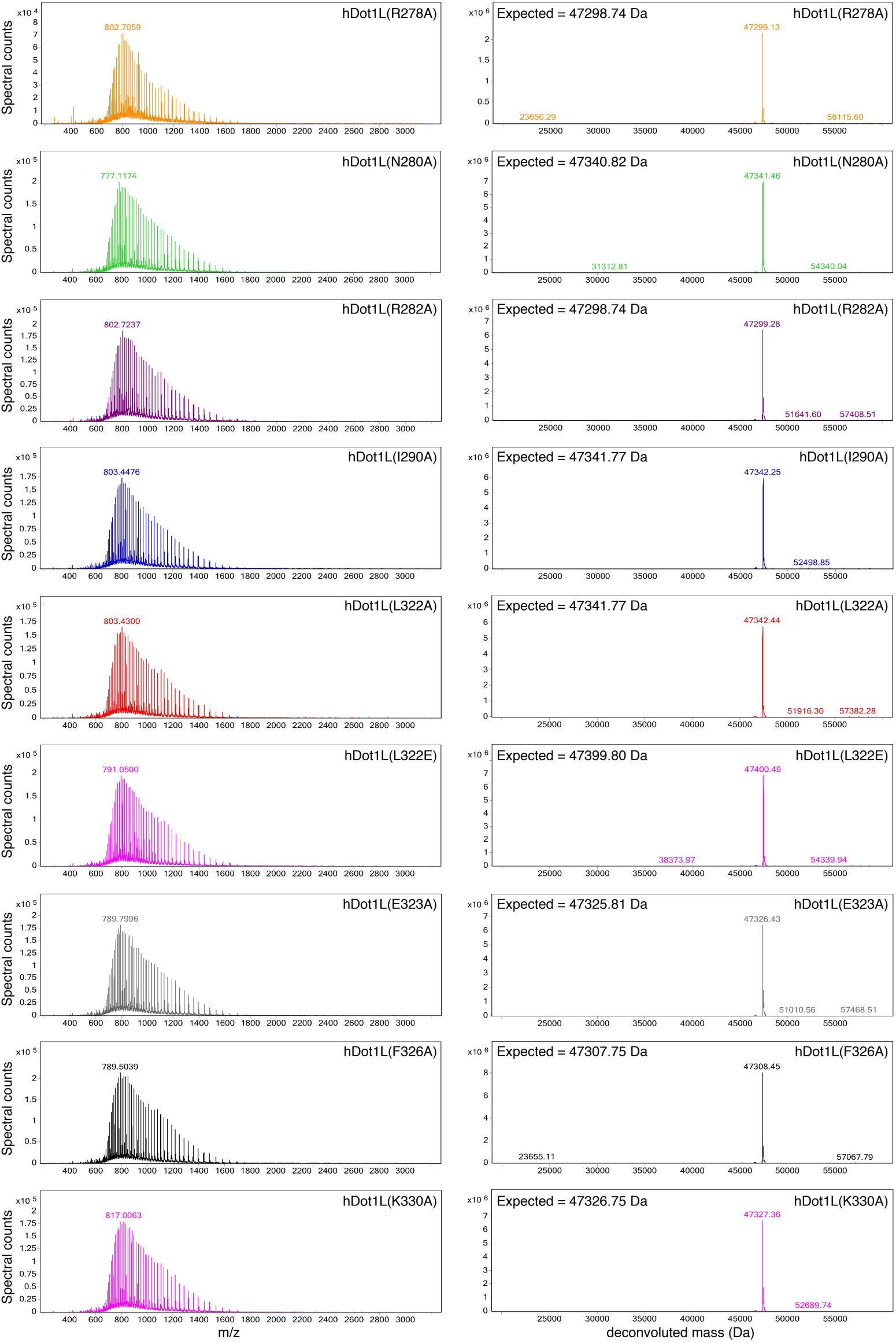
Mass spectrometry characterization of Dot1L mutant proteins. ESI-mass spectra of (left) and deconvoluted mass spectra (right) for mutant Dot1L proteins.

